# Propofol-mediated loss of consciousness disrupts predictive routing and local field phase modulation of neural activity

**DOI:** 10.1101/2023.09.02.555990

**Authors:** Yihan (Sophy) Xiong, Jacob A. Donoghue, Mikael Lundqvist, Meredith Mahnke, Alex James Major, Emery N. Brown, Earl K. Miller, André M. Bastos

**Author notes:** Corresponding Author: André M. Bastos. Equally contributing senior authors (Earl K. Miller, André M. Bastos).

## Abstract

Predictive coding is a fundamental function of the cortex. The predictive routing model proposes a neurophysiological implementation for predictive coding. Predictions are fed back from deep-layer cortex via alpha/beta (8-30Hz) oscillations. They inhibit the gamma (40-100Hz) and spiking that feed sensory inputs forward. Unpredicted inputs arrive in circuits unprepared by alpha/beta, resulting in enhanced gamma and spiking. To test the predictive routing model and its role in consciousness, we collected data from intracranial recordings of macaque monkeys during passive presentation of auditory oddballs (e.g., AAAAB) before and after propofol-mediated loss of consciousness (LOC). In line with the predictive routing model, alpha/beta oscillations in the awake state served to inhibit the processing of predictable stimuli. Propofol-mediated LOC eliminated alpha/beta modulation by a predictable stimulus in sensory cortex and alpha/beta coherence between sensory and frontal areas. As a result, oddball stimuli evoked enhanced gamma power, late (> 200 ms from stimulus onset) period spiking, and superficial layer sinks in sensory cortex. Therefore, auditory cortex was in a disinhibited state during propofol-mediated LOC. However, despite these enhanced feedforward responses in auditory cortex, there was a loss of differential spiking to oddballs in higher order cortex. This may be a consequence of a loss of within-area and inter-area spike-field coupling in the alpha/beta and gamma frequency bands. These results provide strong constraints for current theories of consciousness.

**Significance statement:** Neurophysiology studies have found alpha/beta oscillations (8-30Hz), gamma oscillations (40-100Hz), and spiking activity during cognition. Alpha/beta power has an inverse relationship with gamma power/spiking. This inverse relationship suggests that gamma/spiking are under the inhibitory control of alpha/beta. The predictive routing model hypothesizes that alpha/beta oscillations selectively inhibit (and thereby control) cortical activity that is predictable. We tested whether this inhibitory control is a signature of consciousness. We used multi-area neurophysiology recordings in monkeys presented with tone sequences that varied in predictability. We recorded brain activity as the anesthetic propofol was administered to manipulate consciousness. Compared to conscious processing, propofol-mediated unconsciousness disrupted alpha/beta inhibitory control during predictive processing. This led to a disinhibition of gamma/spiking, consistent with the predictive routing model.

## Introduction

The idea that the brain makes inferences from environmental inputs has been generally accepted ^1^. Predictive coding is a prominent theoretical model of the inferential brain. It argues that previous experiences help construct an internal model which forms feedback predictions, and unexpected stimuli are fed forward, signaling prediction errors ^2–4^. Predictive coding is argued to be involved in cortical functions such as visual ^5,6^ and auditory processing ^7^. Emerging cognitive paradigms and recording techniques have also revealed the potential involvement of predictive coding in higher level cognition such as pattern recognition ^8,9^.

The current paper tests a recent perspective, the predictive routing model, which implements predictive coding in the biological brain by mapping feedforward/feedback streams onto layer-specific brain rhythms and intrinsic anatomical connectivity ^10^. Predictions are fed back via deep-layer cortex alpha/beta (8-30Hz) to prepare cortex for receiving predicted input, and deviations from the predicted inputs (prediction errors) are fed forward via superficial-layer cortex gamma (40-100Hz) and spiking ^11,12^. This model is supported by anatomical and neurophysiological findings. Anatomically, feedback neuronal connections originate in deep layers of higher order cortex and project onto superficial layers of lower order cortex ^13,14^. Evidence from spike-field coherence indicates neurons in deep layers are alpha/beta coherent ^15^. Multi-laminar recordings indicate that alpha/beta rhythms are primarily seen in superficial layers, and local field potential power is primarily seen in deep layers ^16,17^. This can be computationally modeled as alpha-rhythmic inputs to the superficial dendrites of deep layer pyramidal neurons, indicating potential role of L5 Martinotti cells ^18^.

An important theoretical extension from predictive coding to predictive routing is the role of oscillations in carrying out prediction and prediction error computations, instead of single neurons functioning as prediction error computation units. The predictive routing model specifically argues that feedforward gamma/spiking is inhibited by the alpha/beta feedback stream through the respective cortical layer-specific connections ^10^. The current paper aims to causally test this model by measuring putative prediction error responses in the awake state, as well as anesthetized state using propofol, which profoundly decreases alpha/beta oscillations in posterior areas ^19^.

Propofol is a common anesthetic agent that enhances GABAergic activity, inducing loss of consciousness (LOC) ^20^. Under propofol-mediated LOC, the overall neuronal excitability, as measured by single neuron spike rate, is reduced across all cortical areas, and particularly strongly reduced in lateral prefrontal cortex and frontal eye field than in posterior auditory cortex ^19^. In addition, alpha/beta power is strongly reduced in posterior cortex (parietal and visual cortex), as is cortico-cortical as well as cortico-thalamic alpha/beta phase coherence ^19,21^. Previous work has linked interareal alpha/beta coherence to feedback influences ^11,12,14^. Here, we tested how this relatively greater decrease in frontal excitability (compared to sensory cortex excitability), together with a strong reduction in posterior alpha/beta power/coherence, would impact the processing of unexpected stimuli as a model of the predictive routing paradigm. We collected intracranial spike/LFP data in two macaque monkeys before and during propofol-mediated unconsciousness while monkeys were presented with patterns of more or less predictable auditory stimuli. We hypothesized that since alpha/beta-dependent feedback is decreased during propofol-mediated LOC, that feedforward prediction error signals would become disinhibited (therefore increase in signal strength).

Two major models of consciousness have been under scientific discussion as of late: Global Neuronal Workspace theory (GNW) ^22^ and the Integrated Information Theory (IIT) ^23^. GNW posits that the subjective experience of consciousness emerges from communication in the prefrontal-parietal network, and that it is dependent on long-range broadcasting of information in this network. In contrast, IIT takes the approach of formalizing states of consciousness using a metric known as ϕ. It is calculated from binarized neural data (e.g., fMRI or neurophysiology data) and reflects the cause-effect structure present in the signal. From this formalism, proponents of IIT have theorized that a posterior ‘hot zone’ is likely the more important substrate of conscious experience ^24^. In this account, neuronal activation of the ‘hot zone’ is sufficient to induce consciousness experience. Thus, the two theories diverge in their proposed network location of the neural correlates of consciousness, i.e., prefrontal-parietal in GNW and posterior ‘hot zone’ in IIT. Empirical testing of these perspectives has been called upon ^25^ and the current study can inform this debate.

In this study we provide two primary points of evidence to inform the GNW vs. IIT debate. First, we examine which areas (sensory vs. frontal) remain activated by an unpredicted stimulus during propofol-mediated unconsciousness. Under the GNW theory, when subjects are unconscious, posterior areas may remain active but frontal areas will be inactive during processing of unpredictable stimuli. Under IIT, unconsciousness will be primarily reflected in deactivation and/or dis-coordination of posterior cortex. Second, we quantified the within- and inter-areal network coordination by calculating the strength of phasic coupling between alpha/beta and gamma LFP and spiking activity (similar to recent reports, e.g. ^26^). Within-area, spike-phase coupling is thought to reflect the strength of functionally-relevant local computations with increased coupling in gamma and decreased coupling in alpha/beta indexing enhanced computations ^15,27^. Inter-areally, spike-phase coupling is thought to reflect the strength of interareal communication ^28,29^. With respect to network coordination, GNW hypothesizes that unconsciousness will primarily involve the loss of network coordination involving frontal cortex. IIT on the other hand hypothesizes that a loss of network coordination within the posterior ‘hot zone’ will primarily explain LOC.

In the current experiment, monkeys undergo a passive auditory oddball paradigm to elicit neuronal prediction and prediction error signals during propofol-mediated unconsciousness. The auditory oddball paradigm has been used to generate prediction error signals in humans and non-human primates ^8,9,30^. A series of five pure tones is presented, with the first four always repeating (generating prediction), and the last tone having a different frequency (generating prediction error), which is referred to as local oddball (*Figure 1a*). Note that the paradigm also includes global oddballs, which are sequences of repeated stimuli that are rare. However, we did not find significant global oddball responses in this dataset (*Supplemental Figure 1*), and therefore restrict our analyses to local oddballs only. In this report, we will refer to local oddballs simply as “oddballs”. This passive auditory paradigm allows us to investigate neural coding differences both under awake state and following LOC when feedback alpha/beta is disrupted.

**Figure 1.**
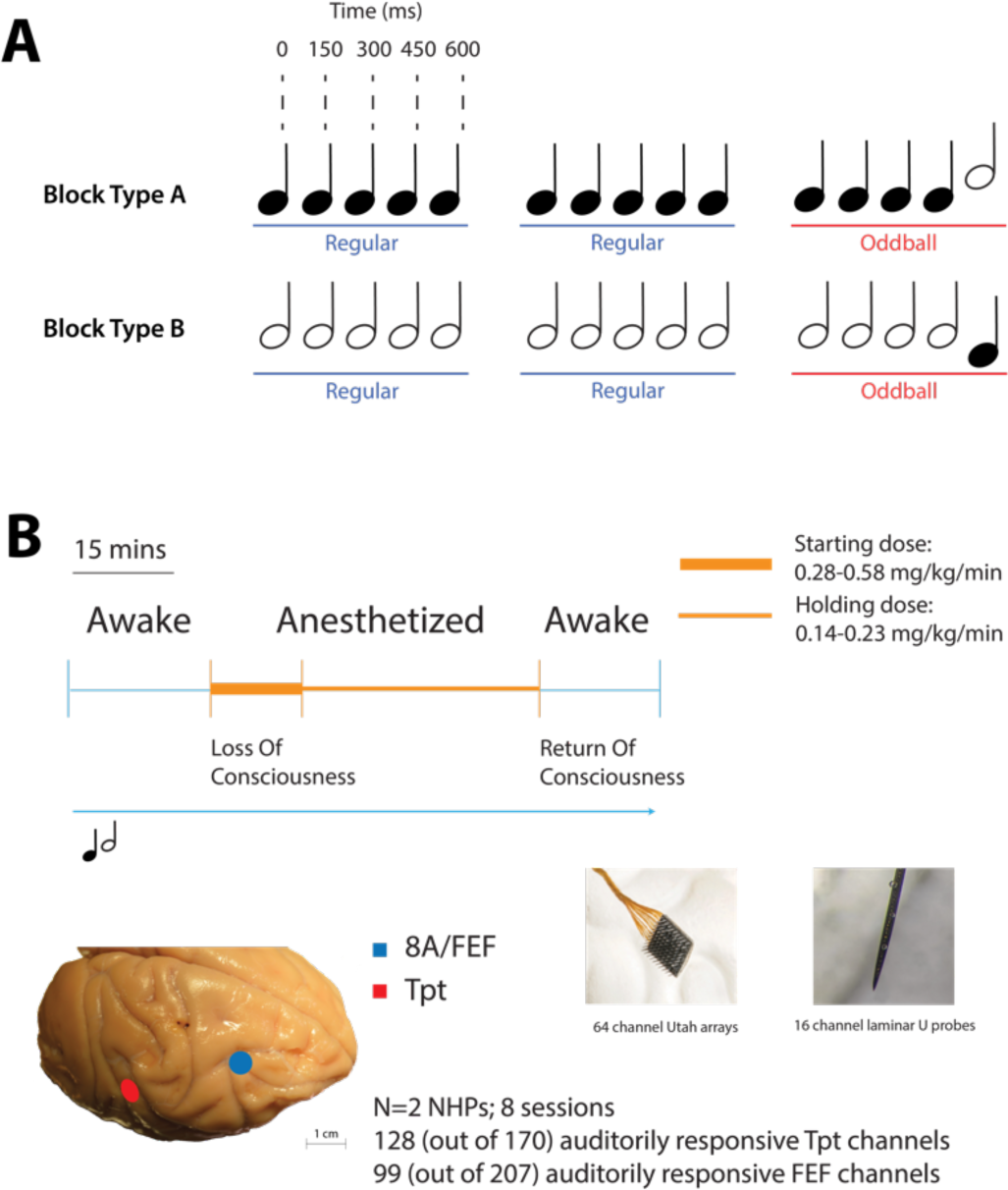
Panel A: Passive auditory oddball paradigm. Series of five tones are presented passively. Each block contains two types of patterns (AAAAA/BBBBB, “regular trials” vs. AAAAB/BBBBA, “oddball trials”). Panel B: propofol infusion was initiated at a higher dose for 20 min and maintained at a lower dose for 40 min. Two types of probes were use (64-channel Utah arrays and 16-channel laminar U-probes). Neural data from areas Tpt and FEF were recorded..

To investigate the hierarchical changes of predictive processing in the cortex, neural activity was recorded at various levels of the cortical hierarchy. This paper primarily focuses on the differential predictive effects in area Tpt (temporoparietal area, an auditory-responsive area in superior temporal gyrus) and area FEF (frontal eye fields, a higher-order frontal area). To note, these two areas have been established to be anatomically connected ^31^. Hierarchical representation of prediction and prediction error has been proposed in the human cortex with data from non-invasive recording methods as well as intracranial recordings ^32,33^. Thus, the current paper expands on this prior work by combining multi-area intracranial data from non-human primates including spiking, LFP, and cortical layer information, with a disruption of alpha/beta connectivity and consciousness via the anesthetic propofol (*Figure 1b*).

## Results

### Disinhibition of feedforward gamma and elimination of feedback alpha/beta modulation during oddball processing

It has been established that unpredictable stimuli evoke an increase in gamma-frequency power and a decrease in alpha/beta low frequency power in both sensory and higher-order cortex ^9,10,32,34,35^. Previous work has also shown a strong reduction in Tpt alpha/beta power and a reduction in fronto-parietal alpha/beta phase coherence during propofol ^19,21^. In predictive routing, alpha/beta oscillations inhibit gamma. Therefore, we hypothesized that during propofol-mediated unconsciousness, cortex would be in an uninhibited state following an oddball stimulus.

To test this, we first compared LFP power in the awake state during oddball trials versus regular (non-oddball) trials. Out of a total of 207 channels recorded in FEF and 170 channels recorded in Tpt, we limited our analysis to 99 auditory-responsive channels in FEF and 128 auditory-responsive channels in Tpt, see Methods. Consistent with predictive routing, both area Tpt and FEF showed increased gamma frequency (> 60 Hz) power following an oddball, and increased alpha/beta frequency (15-30Hz) power following a non-oddball (*Figure 2A, B, upper subpanels*). We then compared oddball versus regular trial LFP power during unconsciousness. Gamma power increased in both area Tpt and FEF during oddball (*Figure 2A, B, lower subpanels*). In area Tpt, the gamma power percent increase to oddball during unconsciousness was higher than the awake state, suggesting disinhibition of the feedforward stream. The center frequency of the gamma power peak during 200ms after oddball onset lowered from 82 Hz during awake state to 40 Hz during unconsciousness. Notably, alpha/beta decrease to oddballs was lost during unconsciousness in both areas (*p* < .05, *Figure 2C*). This is consistent with the hypothesized inhibitory role of alpha/beta in predictive routing, leading to disinhibition of incoming sensory feedforward gamma from lower cortical areas.

**Figure 2.**
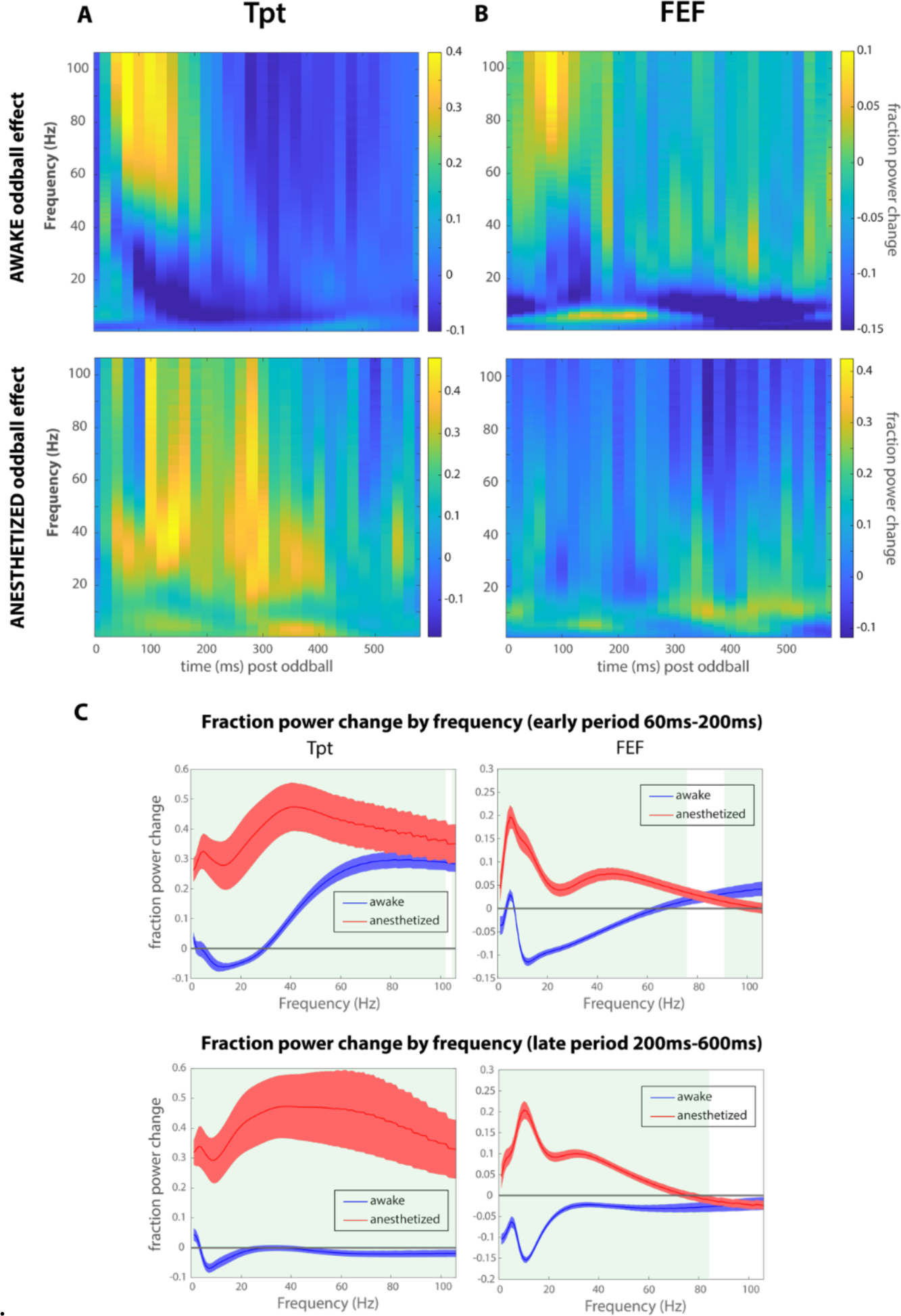
Panel A-B: Fraction power change from regular to oddball trials. Warmer color represents higher power in oddball trials, and cooler color represents higher power in regular trials. Panel A shows results from awake state and Panel B shows results from during unconsciousness. Panel C: Fraction power change (oddball > regular) by frequency for early vs. late period. Shading shows 2*SEM. Green blocks represent time periods during which the awake vs. propofol power change is significantly different.

### Loss of differential spiking to oddball in frontal cortex (FEF) but not sensory cortex (Tpt) during unconsciousness

The predictive routing model hypothesizes that as stimuli become predictable, neural responses are inhibited. Here, we use stimulus temporal repetition as a proxy for low-level stimulus predictability. Note that this equates prediction error responses with release from neural adaptation. However, we primarily examine the differences in prediction error responses as a function of different conscious states wherein low-level adaptation mechanisms should be preserved ^36^. We hypothesize decreased spiking to each repeated stimulus presentation in the regular trial sequence, as well as the first three repeating tones in the oddball sequence (*Figure 1A*). When unpredicted stimuli are presented, the model predicts that neural activity will increase. Since non-sensory related (e.g., baseline spiking) higher order area spiking is known to be more strongly reduced under propofol-mediated unconsciousness than lower order area spiking ^19^, we hypothesize that increased spiking to oddballs in higher order cortex (FEF) will be more strongly reduced during unconsciousness than that in lower order cortex (Tpt).

As expected, multi-unit activity (MUA) decreased throughout the initial 4 tones during all trials (regardless of trial type) both in area Tpt and FEF during awake state (*Figure 3*). MUA response increased to oddball tones during awake state both in area Tpt and FEF (*Figure 3A,B*). During propofol-mediated unconsciousness, a late differential response component was preserved in area Tpt, but there is no strong initial (< 100 ms) peak (*Figure 3C*). In contrast, FEF differential response to oddball was largely eliminated during unconsciousness (*p* < .05, *Figure 3D*).

**Figure 3.**
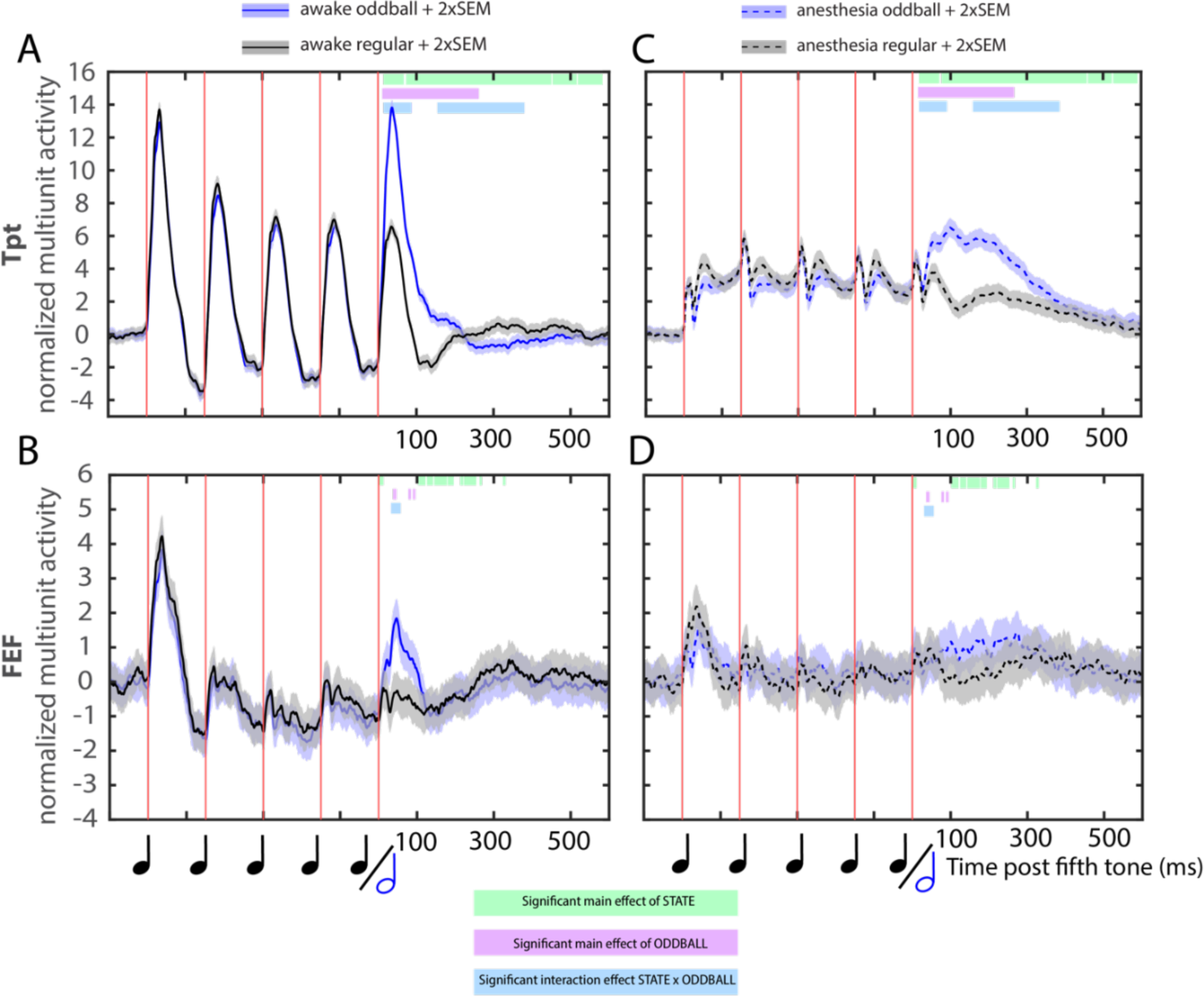
Normalized and baselined multi-unit activity during a 5-tone trial (shading represents ±2*SEM between trials). Red vertical lines show times of tone onset. Horizontal color bars post fifth tone show time clusters of significance (α = 0.05) of main effect of state (awake vs. propofol), main effect of condition (oddball vs. regular), and state by condition interaction. Significant clusters were family-wise error corrected at α = 0.05. N = 128 auditory-responsive Tpt channels, N = 99 auditory-responsive FEF channels. Error bars denote mean +/− 2*SEM across trials.

ANOVA tests were performed at each time point (for every ms), revealing significant clusters in time of main effect of state (awake vs. propofol), main effect of condition (oddball vs. regular), and state by condition interaction. To correct for multiple comparisons, time clusters were family-wise error (FWE) corrected with a non-parametric permutation test at α = 0.05 (see Methods). The results showed a robust state effect of sustained reduction of MUA during propofol-mediated unconsciousness compared to awake state in area Tpt and FEF. Main effect of oddball condition was significant and sustained in time in area Tpt (16-588 ms, with non-significant breaks at 70-74 ms, 456-457 ms, and 518-524 ms) but was significant yet sparse in FEF. The state-by-oddball interaction was significant in both area Tpt and FEF, with Tpt having an early (17-88 ms) reduction and late (158-383 ms) enhancement component and FEF having only one transient component. In sum, these results showed that in Tpt, the late (158-383 ms) spiking increase emerged during unconsciousness compared to awake state. In comparison, oddball responses in FEF were entirely lost during unconsciousness compared to awake state. This suggests that top-down feedback is lost during unconsciousness due to lack of differential spiking to oddballs in frontal cortex, leading to disinhibition of bottom-up input under unconsciousness which allows the late increase component to emerge in sensory cortex.

### Emergence of stronger sink in superficial layers during oddball under unconsciousness, suggesting disinhibition of feedforward stream in microcircuit

Current source density analysis takes the second spatial derivative of the LFP and provides an estimate of laminar current flow in time ^37^. Middle layer inputs (i.e., current sinks in layer 4) are related to feedforward signals from lower-order areas, and superficial layer (layers 2/3) sinks denote feedforward processes that will be sent to higher-order areas ^2,38^. Current sinks were stronger in the middle layer (*Figure 4C*) during oddball compared to regular trials in area Tpt in the awake state (*p* < .05, FWE-corrected at *α = 0.05*, *Figure 4A-C*, note enhanced sink at times 0-200ms post oddball onset). During unconsciousness, this middle layer sink to oddball was eliminated (*Figure 4D*). Instead, a weaker superficial layer sink emerged (*Figure 4F*) and was stronger during oddball than during regular trials (*p* < .05, FWE-corrected at *α = 0.05*, *Figure 4D-F*). ANOVA on the time-space domain with cluster-based permutation test FWE correction (Methods) confirmed the statistical significance of the state by condition interaction (*Figure 4G*), such that the superficial layer source to oddball inverses into a sink during unconsciousness (*Figure 4H*). The emergence of a stronger sink in superficial layers further corroborates with previous evidence in LFP/MUA that the feedforward stream becomes disinhibited during unconscious processing of oddballs.

**Figure 4.**
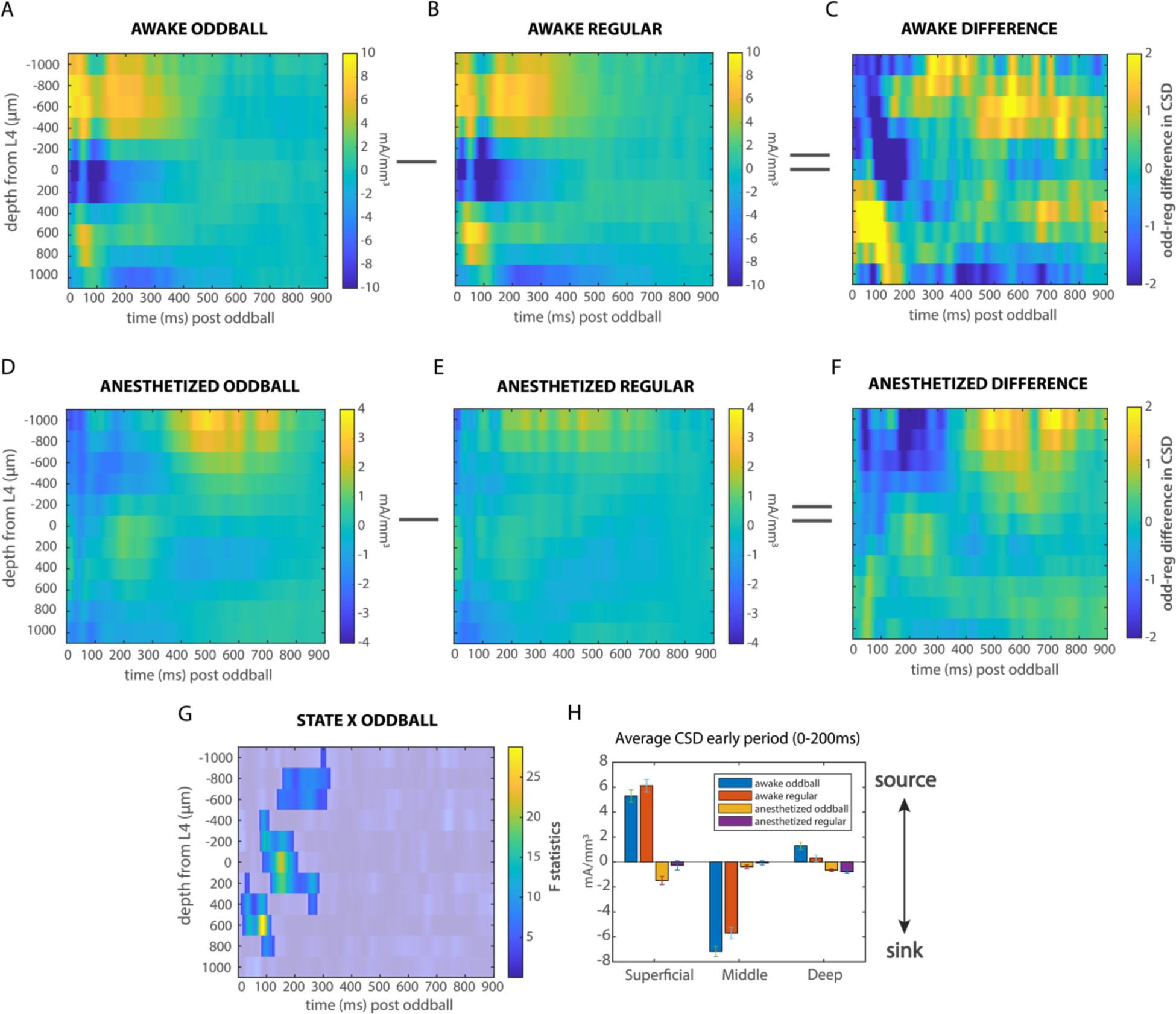
Panel A-F: Sensory region Tpt CSD responses during oddball (panel A) and regular trials (panel B) during awake state; CSD responses during oddball (panel D) and regular trials (panel E) during unconsciousness. Zeros on x-axis indicate the time of oddball presentation. Y-axes represent cortical depth (higher on the graph corresponds to closer to pia mater; negative numbers correspond to superficial layers, e.g. layers 2/3 and positive numbers correspond to deep layers, e.g. layers 5/6. The position of layer 4 is depth 0, see Methods). Difference in CSD between oddball trials vs regular trials in Tpt during awake state (panel C) and during unconsciousness (panel F); Panel G: significant clusters of state-by-oddball interaction effects, FWE-corrected; Panel H: CSD values for superficial vs. middle vs. deep layers for early period (0-200 ms)

### LFP phase – MUA coupling patterns suggested disruption in connectivity following LOC

Slow wave (∼1 Hz) spike-phase coupling is enhanced under propofol-mediated unconsciousness ^19^, which was argued to result in the disruption of spike-phase coupling in faster frequencies such as alpha/beta and gamma ^21^. Since spike-phase coupling in alpha/beta and gamma is considered a mechanism of cortical communication ^15,21,28,39^, the disruption of alpha/beta and gamma synchrony is hypothesized to be a mechanism of anesthesia-mediated LOC. So far, we have demonstrated that during propofol-mediated LOC, oddball responses in sensory cortex are disinhibited relative to the awake state. This was evidenced in area Tpt by a marked increase in gamma power to oddballs, enhanced late (158-383 ms) spiking response to oddballs, and enhanced superficial-layer CSD sink to oddballs. At the same time, spiking and power responses in FEF to oddballs are either diminished or eliminated during propofol-mediated unconsciousness (*Figures 2 and 3*). This begets the question of whether the eliminated FEF responses are due to disruptions in cortical communication either within- or inter-areas (or both).

To quantify neuronal communication, we used a novel metric of spike-phase coupling (see Methods). With this method, the phase of an oscillation is used to fit MUA activity using a sinusoidal basis function. The R^2^ of the fit reveals whether there is a relationship between phase and spiking. The amplitude of the resulting fit is a metric for how powerfully spiking activity is modulated by phase. The LFP phase at which the MUA activity peaks reflects coordination within and across areas, as these interactions may depend on an “optimal” phase ^40,41^. We performed these fits as a function of area, frequency, and conscious state. For within-area spike-phase coupling, sinusoidal functions were fitted to the MUA on each channel as a function of local LFP alpha/beta and gamma phase (mean awake alpha/beta band fit r^2^ = 0.76 in Tpt, r^2^ = 0.77 in FEF; mean awake gamma band fit r^2^ = 0.79 in Tpt, r^2^ = 0.75 in FEF). In alpha/beta band phase to MUA pairs (N = 128 auditory-responsive Tpt channels, N = 99 auditory-responsive FEF channels) the average amplitude of such fits was significantly decreased during unconsciousness (*Figure 5, upper subpanels*, mean awake A = 0.0176 in Tpt, mean A = 0.0081 in FEF; mean propofol A = 0.0055 in Tpt, mean A = 0.0043 in FEF; mean per channel percent decrease in Tpt = 65.6%, mean per channel percent decrease in FEF = 28.6%; rank sum test, *p* < .001), suggesting that the strength of alpha/beta-spiking coupling was significantly weaker during unconsciousness. Gamma band oscillations followed a similar pattern in terms of local modulation of MUA. Propofol-mediated LOC significantly decreased the amplitude of the sinusoidal fits on local MUA as a function of LFP (*Figure 5, upper subpanels*, mean awake A = 0.0220 in Tpt, mean A = 0.0105 in FEF; mean propofol A = 0.0033 in Tpt, mean A = 0.0018 in FEF; mean per channel percent decrease in Tpt = 81.1%, mean per channel percent decrease in FEF = 79.9%; *p* < .001). Interestingly, in area Tpt, gamma coupling was stronger than beta (*Figure 5,* rank sum test, *p* < .05). In area FEF, alpha/beta coupling was stronger than gamma (*Figure 5*, rank sum test, *p* < .05).

**Figure 5.**
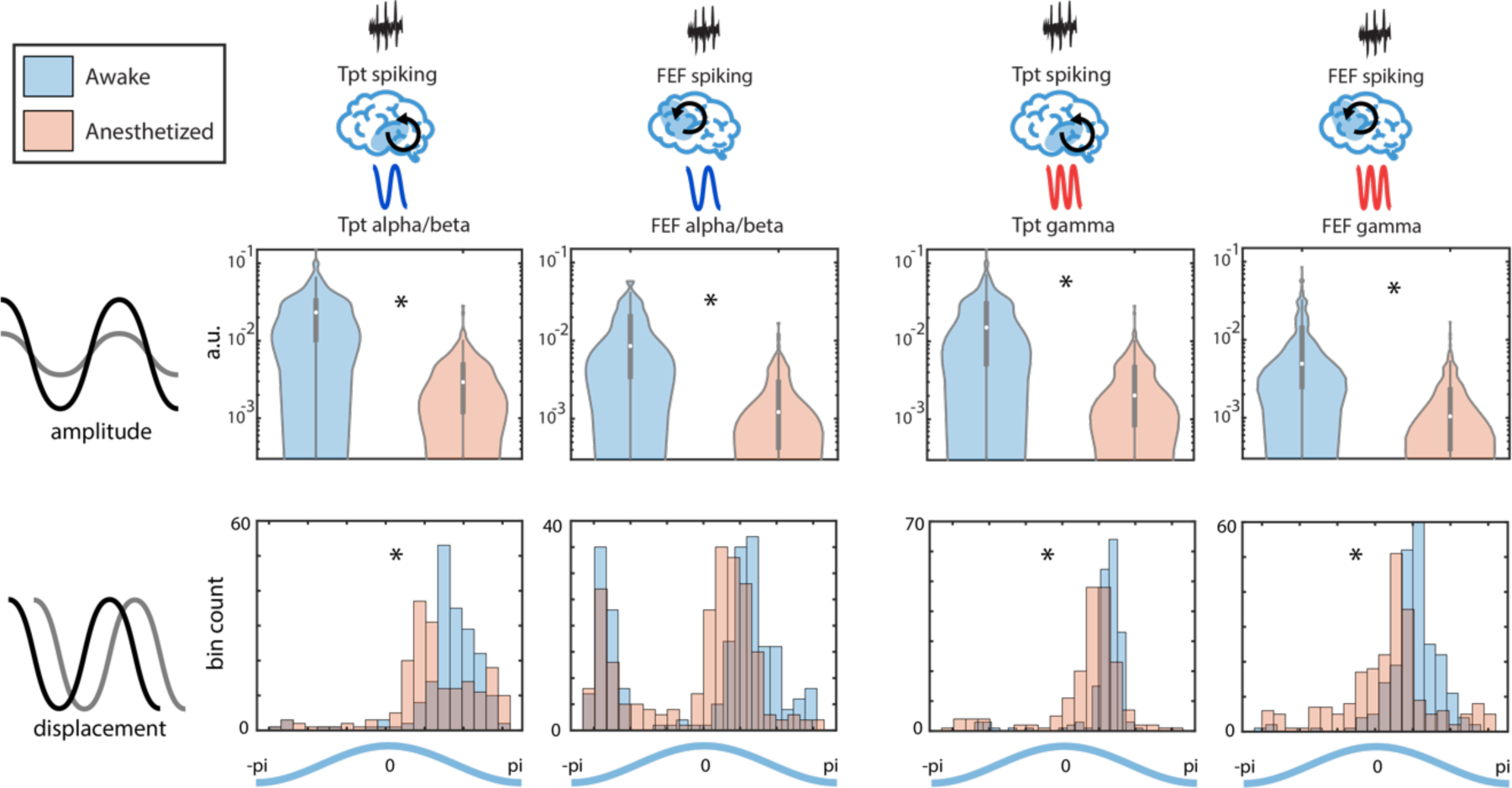
Amplitude and displacement parameters of curve fits to within-area spiking-LFP coupling. N = 128 auditory-responsive Tpt channels, N = 99 auditory-responsive FEF channels. Asterisks (*) indicate significant differences between awake and unconscious states. Violin plot schematics: white dots indicate medians, shaded areas represent kernel density estimates, notches represent upper and lower quartiles, and whiskers indicate range.

The phase relationship between incoming spikes and the phase of an oscillation has been causally shown to modulate neuronal communication and behavioral responses ^41,42^. What this prior work demonstrated is that the phase of incoming spikes may critically determine their impact on the local network. To investigate the effect of propofol on such phase specificity of spike-phase coupling, we examined the displacement parameters of the sinusoidal fits, which corresponds to the peak-to-trough phase specificity of the spike-field coupling. Propofol significantly shifted and disrupted this specificity in both frequency bands (*Figure 5, lower subpanels*). In alpha/beta band, LFP spike-phase coupling had a mean displacement of 1.62 rad (peak to trough) in Tpt in awake state, compared to 1.26 rad (between peak to trough, but phase advanced ←) during unconsciousness (rank sum test, *p* < .05), and 0.02 rad (around peak) in FEF, compared to −0.15 rad (around peak) during unconsciousness (*ns*). In gamma band, LFP phase coupling with spiking has a mean displacement of 1.15 rad (peak to trough) in Tpt in awake state, compared to 0.71 (peak to trough, but phase advanced ←) rad during unconsciousness (*p* < .05), and 1.00 rad (peak to trough) in FEF, compared to 0.35 (around peak, phase advanced ←) rad during unconsciousness (rank sum test, *p* < .05). Of note, the distribution of FEF spike-alpha/beta coupling displacement was bimodal, unlike spike-phase coupling in other frequency/areas.

To test the effect of propofol on interareal LFP spike-phase coupling, the same sinusoidal fitting procedure was done for all interareal channel pairs (Tpt phase - FEF MUA, and FEF phase - Tpt MUA, for both alpha/beta and gamma bands). Overall, the interareal goodness-of-fit was lower than that of local (but still significant) sinusoidal fits in both frequency bands (alpha/beta: mean awake r^2^ = 0.59 from Tpt MUA - FEF LFP phase fits, mean awake r^2^ = 0.55 from FEF MUA - Tpt LFP phase fits; gamma: mean awake r^2^ = 0.59 from Tpt MUA - FEF LFP phase fits, mean awake r^2^ = 0.47 from FEF MUA - Tpt LFP phase fits). Consistently, the amplitudes of the sinusoidal fits were significantly decreased in both interareal directions and both frequency bands (*Figure 6, upper subpanels*, alpha/beta: mean A = 0.0022 for Tpt MUA by FEF LFP phase fits during awake state, mean A = 0.0014 during unconsciousness, mean per channel percent decrease = 24.8%; mean A = 0.0013 for FEF MUA by Tpt LFP phase fits during awake state, mean A < 0.0001 during unconsciousness, mean per channel percent decrease = 31.9%; gamma: mean A = 0.0016 from Tpt MUA by FEF LFP phase during awake state, mean A < 0.0001 during unconsciousness, mean per channel percent decrease = 58.3%; mean A = 0.0010 from FEF MUA by Tpt LFP phase during awake state, mean A < 0.0001 during unconsciousness, mean per channel percent decrease = 61.3%).

**Figure 6.**
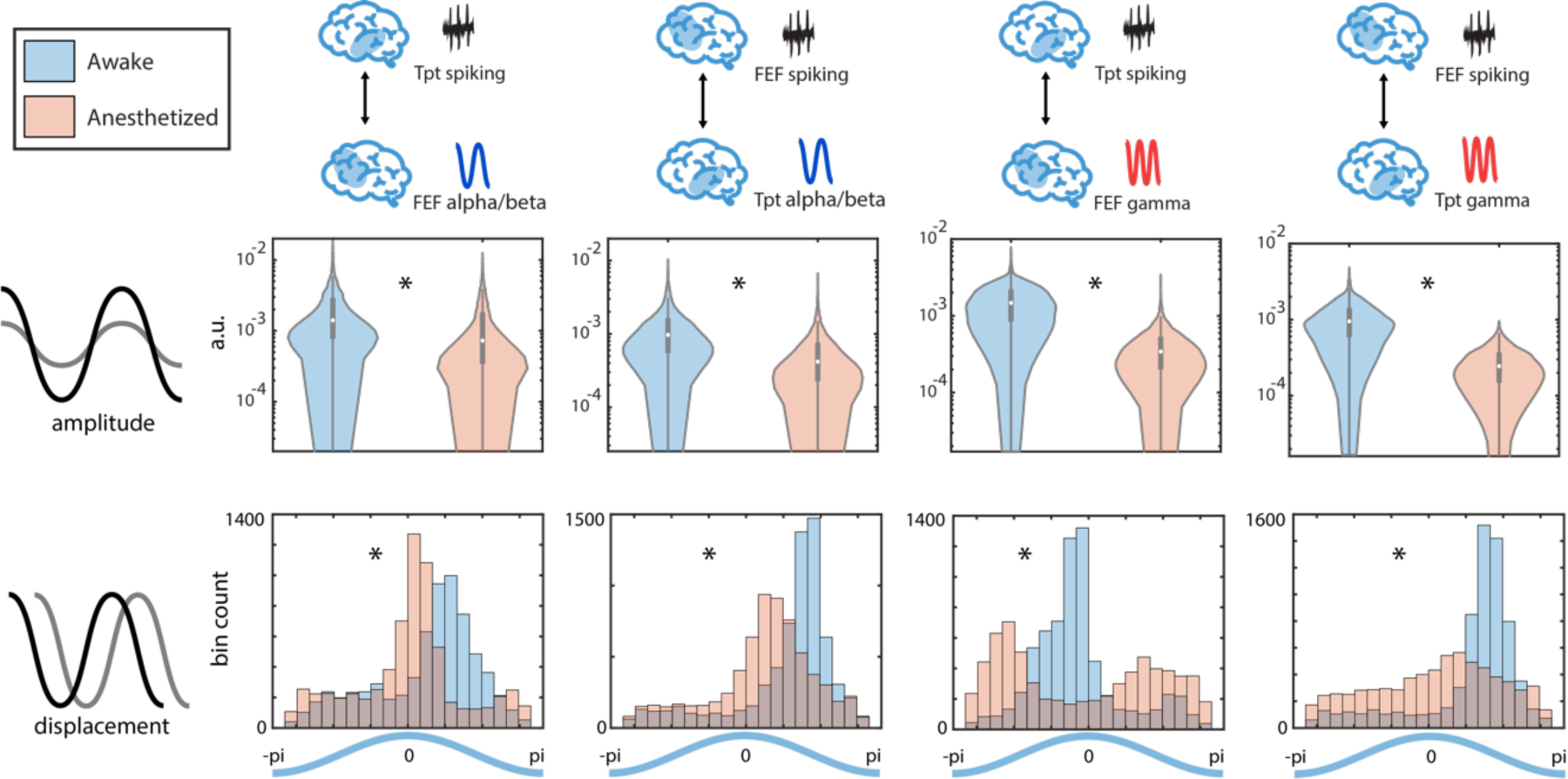
Amplitude and displacement parameters of curve fits to interareal spiking-LFP coupling. N = 128 auditory-responsive Tpt channels, N = 99 auditory-responsive FEF channels. Asterisks (*) indicate significant differences between awake and unconscious states. Violin plot schematics: white dots indicate medians, shaded areas represent kernel density estimates, notches represent upper and lower quartiles, and whiskers indicate range.

Sinusoidal fits of interareal LFP phase-spiking coupling also showed displacement shifts during unconsciousness (*Figure 6, lower subpanels*). In alpha/beta band, Tpt LFP phase coupling with FEF MUA had a mean displacement of 0.50 rad (peak to trough) in awake state, compared to 0.03 rad (around peak, phase advanced ←) during unconsciousness (rank sum test, *p* < .05). FEF LFP phase coupling with Tpt MUA had a mean displacement of 1.07 rad (peak to trough) in awake state, compared to 0.47 rad (peak to trough, phase advanced ←) during unconsciousness (rank sum test, *p* < .05). In gamma band, Tpt LFP phase coupling with FEF MUA had a mean displacement of −0.30 rad (trough to peak) in awake state, compared to −0.21 rad (trough to peak, but phase delayed →) during unconsciousness (rank sum test, *p* < .05); FEF LFP phase coupling with Tpt MUA had a mean displacement of 1.12 rad (peak to trough) in awake state, compared to 0.20 rad (peak to trough, but phase advanced) during unconsciousness (rank sum test, *p* < .05).

## Discussion

### Interruption of the inhibitory feedback component of predictive routing in LFP, MUA, and CSD

To test the predictive routing model and its role in conscious processing, we devised this study to include both feedforward (prediction error elicited by oddball trials) and feedback (prediction established by repeated regular trials) components, under awake state vs. propofol-mediated LOC. Cortical feedforward information has been proposed to be carried by gamma and feedback by alpha/beta ^11,12,14,43^. The predictive routing model posits that alpha/beta band prepares sensory cortex by inhibiting representations that are predicted ^10^. This was corroborated by our results in the awake state, in which gamma power was increased to oddball trials and alpha/beta power was decreased. Many have argued that the feedback component is an integral information stream to support consciousness ^22,44^. In line with the predictive routing model, the absence or degradation of feedback stream will disrupt predictive processing as well as the state of consciousness.

In sum, we observed a disruption of the feedback stream during unconsciousness. In LFP power, propofol-mediated LOC significantly reduced alpha/beta power during prediction, both in auditory and frontal cortex. Frontal cortex spiking to oddballs were eliminated during unconsciousness. In area Tpt, there was a significant loss of spiking in the early period to oddballs during unconsciousness. Loss of initial early peak is suggestive of decreased feedforward inputs from earlier sensory areas ^45^, likely reflecting the global GABAergic effect of propofol. Next, there was an enhanced and sustained oddball (prediction) spiking response in sensory area Tpt. This delayed enhancement is in line with the hypothesis that inhibitory feedback from higher cortex is disrupted. This disruption of the feedback stream allows the feedforward stream to become disinhibited. This is also exhibited in the prolonged enhancement of prediction error gamma power during unconsciousness (compared to awake state, Fig. 2A). Similarly, the emergence of a greater superficial sink during prediction error vs. prediction also suggests the disinhibited feedforward process hypothesis. This greater superficial sink is well-positioned near L3, which is the known output layer for feedforward projections ^46^.

Also of note is the hierarchical nature of oddball in multi-unit activity. Prediction error processing was preserved (and prolonged) only in auditory cortex under propofol-mediated LOC. In FEF, differential response to prediction error in MUA is eliminated. This hierarchical difference also suggests that predictive processing is not merely a result of neuronal adaptation, exhibited as decrease in firing rate due to local cellular mechanisms ^47^. For example, the strength of neuronal adaptation also shows hierarchical differences. FEF spiking adapts to near-baseline after a single auditory tone, whereas Tpt spiking slowly adapts over several tones.

### Phasic coupling of LFP and MUA within- and inter-areas is disrupted by propofol

Within-area spike-field coupling is often interpreted as the strength of local computation, often functionally relevant in domains such as attention ^15,28,39^. Interareal spike-field coupling, on the other hand, is interpreted as the strength of communication and information flow ^28,29^ (but see ref ^48^). The disruption of local computation and interareal connectivity has been hypothesized to be a mechanism of anesthesia-mediated LOC ^19,49^. In our analysis, we examined the amplitude and phasic shift of a sinusoidal fit from alpha/beta and gamma band phase onto multi-unit activity, both within-area and inter-areas.

Our results reveal that both alpha/beta and gamma band LFP phase strongly modulate both local and interareal multi-unit activity in the awake state. Under propofol-mediated LOC, this phasic modulation decreases, both within-area and inter-areas. This suggest that propofol disrupts local computation as well as interareal communication. The phase specificity of within-area spike timing to LFP oscillations has been argued to encourage efficient information encoding via summation of potentials ^50,51^. In other words, there exists an optimal LFP phase to spiking activity peak that allows for maximal information encoding in the cortex ^41,42^. This optimal phase to spiking activity peak is not achieved under propofol, suggesting a lag in communication between neural rhythms caused by this anesthetic.

### Implication on theories of consciousness: IIT vs. GNW?

Recent ongoing debates between two leading theories of consciousness have called for empirical evidence ^25^. The authors list three diverging predictions from each theory. We address two of these predictions with our results below:

#### Prediction #1: location of conscious content

IIT posits that posterior ‘hot zones’ are primarily responsible for conscious content. GNW theory posits that frontal areas (and their connections) are necessary for conscious content. During propofol-mediated unconsciousness, we observed enhanced late spiking, gamma power, and superficial layer current sinks to unpredictable vs. predictable stimuli in Tpt. Our results show that posterior region activity alone is insufficient for conscious perception. Propofol-mediated LOC was marked by the elimination of prefrontal differential spiking to oddballs. Our results therefore suggest an important role for prefrontal cortex activation, in addition to sensory cortex activation, for conscious perception consistent with GNW theory.

#### Prediction #2: interareal connectivity during conscious perception

IIT posits that conscious perception is marked by short-range connectivity in posterior regions. GNW theory posits that conscious perception is marked by long-range connectivity between lower and higher order cortical areas. Our results from spike-phase coupling show that both within-area and interareal connectivity are significantly reduced under LOC. Thus, our results provide evidence for both theories.

In sum, in line with the predictive routing model, alpha/beta oscillations in the awake state serve to inhibit the processing of predictable stimuli. Propofol-mediated LOC eliminates alpha/beta modulation in sensory cortex and alpha/beta phasic modulation between sensory and frontal areas. As a result, we propose, oddball stimuli cause enhanced gamma power, late period spiking, and superficial layer sinks in sensory cortex. Therefore, somewhat paradoxically, auditory cortex is in a disinhibited state during propofol-mediated LOC when processing oddballs. However, despite these enhanced feedforward responses in auditory cortex, there is a loss of an oddball response in higher order cortex. The inability of sensory cortex to effectively drive higher-order cortex may be the result of a loss of within- and inter-areal spike-field coupling in the alpha/beta and gamma frequencies. In addition, these results provide empirical evidence for the IIT vs. GNW debate: activation of posterior sensory cortex is insufficient for conscious processing. Instead, conscious processing is promoted by activation of both sensory and frontal cortices to an oddball, together with intact within- and inter-area oscillatory dynamics at both alpha/beta and gamma frequencies.

## Materials and Methods

(Some text below are recycled from ref ^19^ in cases where methods are identical)

### Experimental Subjects and Propofol Infusion

Two rhesus macaques (*Macaca mulatta*) aged 12 years (monkey 1, male, ∼13.0 kg) and 13 years (monkey 2, female, ∼9.5 kg) participated in the study. Monkey 1 was surgically implanted with a subcutaneous vascular access port (Swirl port mini, Norfolk Access Technologies, Skokie, IL) at the cervicothoracic junction of the neck with the catheter tip reaching the termination of the superior vena cava via the external jugular vein. Monkey 2 was acutely implanted with a 24g Terumo surflo catheter to a vein in the ear for each session. Propofol infusion was initiated at a higher dose (285 µg/kg/min for monkey 1; 285-340 µg/kg/min for monkey 2) for 20 min, and maintained at a lower dose (142.5 µg/kg/min for monkey 1; 143-200 µg/kg/min for monkey 2) for 40 min.

### Electrophysiological Recordings

Neurophysiology with chronic Utah arrays: For recordings in cortex, monkeys 1 was chronically implanted with four 8 x 8 iridium-oxide contact microelectrode arrays (‘Utah arrays’, MultiPort: 1.0 mm shank length, 400 mm spacing, Blackrock Microsystems, Salt Lake City, UT), for a total of 256 electrodes. Arrays were implanted in the prefrontal (area 46 ventral and 8A), posterior parietal (area 7A/7B), and temporal-auditory (caudal parabelt area STG [superior temporal gyrus]) cortices. Specific anatomical targeting utilized structural MRIs of each animal and a macaque reference atlas, as well as visualization of key landmarks on surgical implantation ^52^. For Utah array recordings, area 8A and PFC were ground and referenced to a common subdural site. Area STG and PPC also shared a common ground/reference channel which was also subdural.

To ensure signals recorded on the multiple data acquisition systems remained synchronized with zero offset, a synchronization test signal with locally unique temporal structure was recorded simultaneously on one auxiliary analog channel of each system. The relative timing of this test signal between each system’s recorded datafile was measured offline at regular intervals throughout the entire recording session. Any measured timing offsets between datafiles were corrected by appropriately shifting spike and event code timestamps in time, and by linearly interpolating analog signals to a common time base.

Neurophysiology with acute laminar probes: For recordings in monkeys 2, the monkey was first implanted with a custom-machined Carbon PEEK chamber system with three recording wells placed over visual/temporal, parietal, and frontal cortex ^53^. We acutely introduced 16 or 32 contact ‘multi-laminar’ probes (U/V probes, Plexon, Dallas, TX) into the same cortical areas we recorded with chronic Utah arrays: areas STG, 7A/7B, 8A, and VLPFC. Between 1 and 2 probes were used per recording chamber and a total of 4–6 probes were used per session. The total channel count ranged between 96 and 128 electrodes per session. Electrode contacts on these probes were spaced 100 mm apart for the 32 channel probes or 200 mm apart for the 16 channel probes. This gave a total linear sampling of 3.0–3.1 mm on each probe. The recording reference was the reinforcement tube, which made metallic contact with the entire length of the probe (total probe length from connector to tip was 70 mm). With MRI guidance, we introduced the probes to be perpendicular to cortex and to span all cortical layers, as previously described ^10^. As a marker for layer 4, we used the relative power profiles calculated in the pre-propofol awake state. The cross-over between the relative power of the gamma and alpha-beta bands was used to estimate the location of layer 4 ^17^.

### Experimental Paradigm

Our paradigm is adapted from a previous auditory oddball paradigm design in Bekinschtein et al., 2009. Series of five pure tones of 50 ms were presented. Each five tones (referred to as a trial) consist of two different frequencies (Tone A = 800 Hz, Tone B = 1600 Hz). The inter-tone interval within a trial is 150 ms. The inter-trial interval is fixed to 500 ms. Four different stimulus blocks were used: AAAAA, BBBBB, AAAAB, and BBBBA blocks. In AAAAA blocks, 20 AAAAA sequences were delivered, followed by a random mixture of 80 AAAAA and 20 AAAAB. In BBBBB blocks, 20 BBBBB sequences were delivered, followed by a random mixture of 80 BBBBB and 20 BBBBA. In AAAAB blocks, 20 AAAAB sequences were delivered, followed by a random mixture of 80 AAAAB and 20 AAAAA. In BBBBA blocks, 20 BBBBA sequences were delivered, followed by a random mixture of 80 BBBBA and 20 BBBBB.

### Data Preprocessing

LFPs were recorded at 30 kHz and filtered online via a lowpass 250 Hz software filter and downsampled to 1 kHz. Spiking activity was recorded by sampling the raw analog signal at 30 kHz, bandpass filtered from 250 Hz to 5 kHz, and manually thresholding. Blackrock Cereplex E headstages were utilized for digital recording via 2–3 synchronized Blackrock Cerebus Digital Acquisition systems. Analog multi-unit activity was extracted with band-pass filtering at 300–4500 Hz with a zero-phase fourth-order Butterworth filter. Data was then epoch-ed by trial and baselined by standard error taken from 500ms pre-stimulus onset.

Time frequency analysis was performed with the Fieldtrip toolbox (http://www.fieldtriptoolbox.org/) ^54^ on MATLAB (The Mathworks, Inc, Natick, MA) separately for the frequency bands discussed (alpha/beta: 1-40 Hz; gamma: 40-150 Hz) with multi-taper-method convolution (8 tapers for alpha/beta bands and 20 tapers for gamma band). Auditory-responsive channels were selected for analyses in this manuscript. We define auditory-responsive channels as channels with si gnificant differences between −200 ms to −100 ms before stimulus onset and 0 ms to 100 ms after stimulus onset (Wilcoxon’s rank sum test).

### Current Source Density Analysis

CSD was computed from LFP as a second spatial derivative estimate using this equation (Nicholson and Freeman 1975):

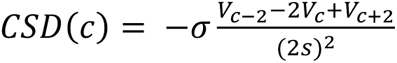

where V refers to the voltage recorded at channel c, s refers to the inter-channel spacing (200 μm) and σ is the tissue conductivity (assumed to be 0.4). Channels were aligned by identifying the earliest sink to stimulus presentation under awake state as input layer. CSD data was then averaged across sessions after laminar alignment (similar to previous work ^16^). CSD output for each trial was Gaussian-filtered with a filter size of [10 1].

### Statistics

Cluster-based significance testing was done based on a nonparametric method for electrophysiology signals outlined in ref ^55^. Two by two (state by trial type) significant clusters (first-level statistics α = 0.05, defined as continuously significant in time for MUA analysis, continuously significant in time and frequency connecting by at least an edge for power spectra, and continuously significant in time and space connecting by at least an edge for CSD profiles) were identified with ANOVAs performed on individual time (for MUA) or time by frequency points (for power spectra and CSD profiles). Trial types were then randomly permuted 1000 times to create a noise distribution of largest significant cluster sizes, and clusters of sizes bigger than 95% of the noise distribution were allowed to survive (i.e. second-level statistics α = 0.05).

### Power binning and MUA correlation

LFP phase was calculated by taking the phase angle between the original signal and its Hilbert transform (using the MATLAB angle() and hilbert() functions) on the raw LFP data. For alpha/beta band phase analysis, the bandpass filter was set to 8 – 35 Hz; for gamma band phase analysis, the bandpass filter was set to 40 – 80Hz. The calculated continuous phase was then divided into 8 bins, and corresponding MUA was taken average per channel according to the phase bin the LFP is in at that timepoint. A one-term Fourier series was then fit to the binned MUA averages using the MATLAB fit() function, which gives:

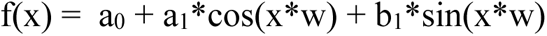

in which w was fixed to be 1 period. Displacements and amplitudes of each fit were then calculated by:

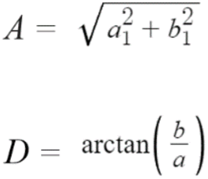

## Acknowledgements

We would like to thank Charles Shvartsman for help with initial analyses of this dataset. This work was supported by NIMH R00MH116100, NIGMS P01GM118269, NIMH R01MH115592, the JPB Foundation, and The Picower Institute for Learning and Memory.

---

**Supplemental Figure 1.**
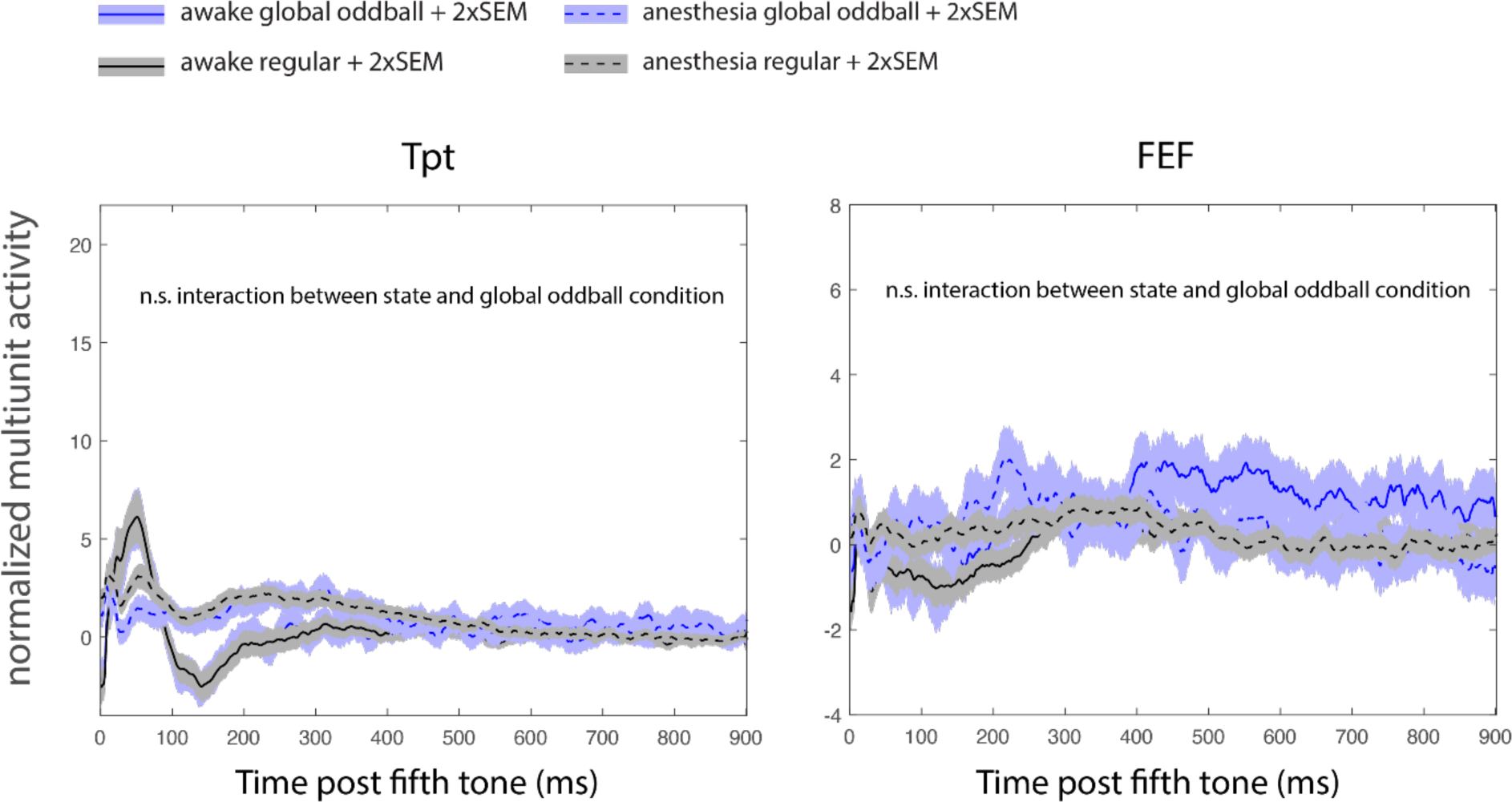
Multi-unit Activity after fifth tone for regular trials vs. “global oddballs” (AAAAA or BBBBB trials that are low probability in a block of high probability of AAAAB or BBBBA)

